# Can digital epidemiology indicate the emergence and current distribution of canine dirofilariosis?

**DOI:** 10.64898/2026.06.16.732585

**Authors:** Tamara Szentiványi, László Zsolt Garamszegi

## Abstract

The emergence and spread of vector-borne pathogens, such as the mosquito-borne parasite *Dirofilaria immitis*, the causative agent of canine heartworm disease pose increasing challenges for public and veterinary health. These parasites have expanded from Southern Europe into Central and Eastern Europe, where locally acquired transmission is increasingly documented in pets and wild animals, and is now considered an emerging infectious disease. Understanding the dynamics of its emergence and public awareness is essential for effective surveillance and intervention. Here, we evaluated whether human online search behavior, as measured by Google Trends (GT), reflects patterns of dirofilariosis emergence in Hungary. We analyzed GT data for heartworm-related terms from 2012 to 2025, exploring geographical and temporal trends. Our analyses revealed a sustained increase in search interest over time beginning in 2012, with pronounced rise after 2015. Spatially, the highest relative search volumes were concentrated in southeastern Hungary. We found a strong positive association between regional search activity and mosquito infection rate. These findings demonstrate that GT-derived data accurately mirror both the temporal increase and geographic distribution of canine dirofilariosis, supporting the use of digital epidemiology as a complementary surveillance tool to inform public health and veterinary strategies for emerging vector-borne diseases.

## Background

The emergence of vectors and vector-borne pathogens present a growing challenge for public health and veterinary medicine [1,2]. Among these, several filarioid nematode species, including *Dirofilaria* spp., have expanded across Europe over the past two decades, alongside their potential vectors, such as invasive mosquito species [3–7]. *Dirofilaria immitis*, the causative agent of heartworm disease, has shifted from being historically endemic in Southern Europe to an increasingly recognized endemic parasite in Central and Eastern Europe, as well [3,8–10]. Across Europe, including Hungary, evidence of autochthonous transmission has become increasingly frequent, since its first detection in 2007, as reflected in veterinary records and molecular surveillance in mosquitoes [10,11,20,21,12–19]. Additionally, an increase in human infection is also observed across Europe [4,22–27]. Nevertheless, its patterns of emergence, geographic expansion, and evolving public awareness are still not fully understood.

In recent years, digital epidemiology and infodemiology have emerged as complementary approaches that use digital data sources to support public health surveillance. Digital epidemiology exploits digital traces, such as internet searches and social media activity, to monitor disease patterns, whereas infodemiology focuses on the distribution of health information and online information-seeking behavior. Together, these fields provide valuable insights into population-level behavior and real-time public interest in health-related topics [28,29]. Tools such as Google Trends (GT), which quantify the relative volume of online search queries, offer a publicly accessible way to monitor interest in symptoms, pathogens, and vector-borne diseases [30–34]. The use of GT as a surveillance proxy is based on the assumption that increases in search activity reflect changes in disease occurrence, parasite abundance, disease impact, or public awareness that prompt individuals to seek information online. However, this assumption has rarely been formally validated and may also be influenced by factors such as media coverage, public health campaigns, or changes in internet use. Despite these limitations, GT has been successfully applied to monitor infectious disease seasonality [35,36], outbreak dynamics [37,38], and public responses to zoonotic and vector-borne threats, including West Nile virus and Lyme borreliosis [33,39–41]. Applications of GT in veterinary parasitology are still relatively limited, though some research has examined trends such as the occurrence of tick paralysis in companion animals [42], the monitoring of swine fever outbreaks [43], and patterns of public interest during canine leishmaniosis outbreaks [44].

Understanding whether human online search behavior reflects real epidemiological trends have practical implications for early warning, targeted communication, and optimizing veterinary resource allocation. Our study investigates whether Google Trends (GT) data can be used to understand the emergence of dirofilariosis in Hungary. First, our goal was to assess the validity of these digital indicators by comparing GT-derived search activity with molecular xenomonitoring data obtained from mosquito populations [19]. Additionally, we aimed to examine temporal and spatial patterns in heartworm-related search queries to determine whether public search activity is consistent with the known occurrence and geographic expansion of the disease. Finally, our goal was to evaluate the potential of online search data, combined with xenomonitoring, as a complementary surveillance tool for monitoring the emergence and distribution of canine dirofilariosis.

## Methods

### Correlations with mosquito xenomonitoring across spatial regions

Spatial patterns of Google Trends (GT) relative search volume (RSV) values for the Hungarian-language terms associated with canine dirofilariosis (“szívféreg” and “szívférgesség”) were compared with previously published molecular xenomonitoring data from mosquito populations [19]. RSV is a normalized index representing the proportion of searches for a given term relative to total Google search activity within a defined geographic region and time period, scaled from 0 to 100, where 100 indicates peak relative search interest. Mosquito infection was quantified at the regional (NUTS-2) level as minimum infection rate (MIR), calculated as the number of *Dirofilaria immitis*-positive mosquito pools per 1,000 mosquitoes tested. To ensure temporal correspondence between the datasets, regional RSV values were averaged over the 2022–2023 period, matching the years during which mosquito collections were conducted. Associations between regional RSV and MIR were assessed using Spearman’s rank correlation coefficient. Because the regional analysis was based on only seven observations and the MIR values were not normally distributed, non-parametric methods were used. This analysis was used to evaluate whether regions with higher mosquito infection rates also exhibited greater public interest in heartworm disease as reflected by online search activity. Mosquito sampling procedures and molecular detection methods have been described previously [19].

### Temporal trends of Google Trends data and disease-specific comparison

GT was used to extract RSV values for Hungarian-language terms associated with canine dirofilariosis (“szívféreg” and “szívférgesség”), along with two control diseases, parvovirus and rabies (“parvovírus” and “veszettség”, respectively) selected based on their relevance in public communication. Searches were restricted to Hungary and extracted for the full available period (2007–2025) to cover the timeframe during which dirofilariosis emergence and spread have been documented. However, analyses were limited to data from 2012 onward, due to minimal search activity in earlier years. All data were obtained using the “search term” option to minimize semantic variability, and RSV values were used without further transformation.

To explore disease-specific temporal trends, monthly Google Trends RSV values from January 2012 to December 2025 were averaged within each calendar year to obtain annual mean RSV values for each search term. These annual mean values were then analyzed using separate linear regression models for each search term, with annual mean RSV as the response variable and calendar year included as a continuous predictor (annual mean RSV ∼ Year). Regression coefficients (β), standard errors (SE), p-values, and coefficients of determination (R²) were used to quantify and compare long-term temporal trends among search terms.

To explore seasonal differences in search activity, monthly RSV values were modeled using a linear modeling framework, with calendar month included as a categorical predictor (12 levels). In complementary analyses, month was included alongside calendar year and search term in a linear model to account for long-term temporal trends and differences among diseases. Statistical significance of seasonal effects was evaluated using analysis of variance (ANOVA).

### Regional and temporal patterns of RSV

Yearly county-level (NUTS-3 regions) RSV values for heartworm-related search terms were also analyzed for the period 2012–2025. Because repeated yearly observations were available for each county, linear mixed-effects models were applied to account for non-independence of measurements within regions. RSV was modeled as a function of calendar year, treated as a continuous predictor. County was included as a random factor to account for non-independence of search intensity due to repeated observations within counties (random intercept model). To assess whether the rate of temporal change differed among counties, a second model including random slopes for year was fitted, allowing the temporal trend to vary by region. Model fit was evaluated using likelihood ratio tests comparing the random intercept and random slope models under maximum likelihood estimation. Additionally, to quantify the degree of clustering of RSV values within counties, repeatability was calculated from the random intercept model based on the estimated variance components. Repeatability represents the proportion of total variance attributable to variance at the among-county level.

All analyses were performed in R Studio version 4.3.1 [45], using the packages tidyverse [46], and lme4 [47]. Figures were generated using packages: ggplot2 [48], gganimate [49]. Maps were created using QGIS version 3.24.1 [50].

## Results

### Correlations with mosquito xenomonitoring across spatial regions

Geographical patterns in RSV exhibited similar spatial variation to what can be observed from minimum infection rate (MIR) (Fig. 1AB) [51]. At the regional (NUTS-2) level, a statistically significant positive association was observed between MIR and RSV, when considering the period mosquitoes were collected in 2022–2023 (Spearman’s ρ = 0.86, p = 0.014) (Fig. 3AB). Similarly, when mean RSV values across the full study period (2012–2025) were considered, regions with consistently higher MIR also exhibited higher long-term search interest (Spearman’s ρ = 0.85, p = 0.013).

**Figure 1AB.**
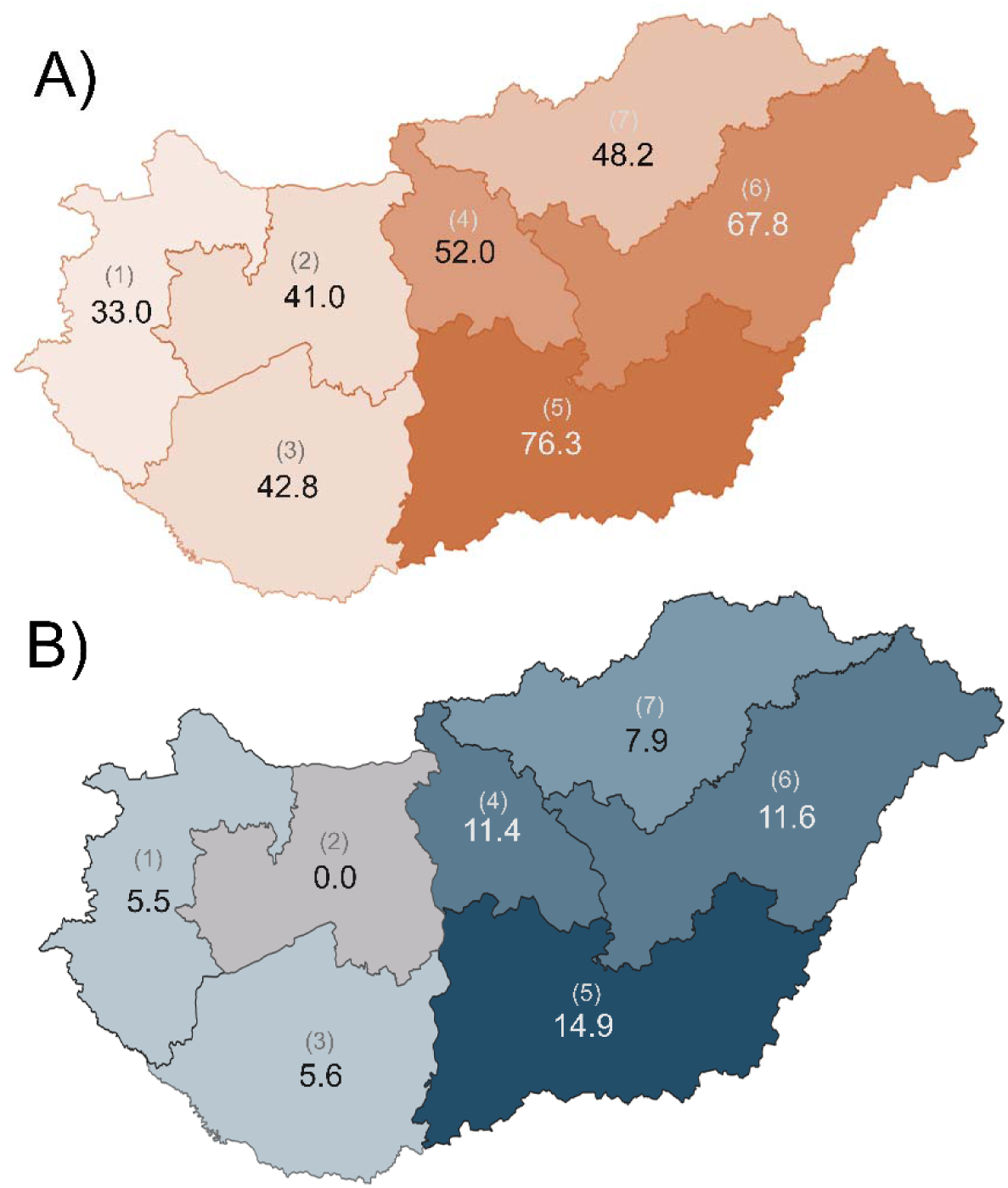
Regional distribution of Google Trends relative search volume (RSV) and mosquito infection rates (MIR). Regional distribution of (RSV) for heartworm-related terms (average of “szívféreg” and “szívférgesség”) for the period 2012–2025 (A); and geographic distribution of *Dirofilaria immitis* minimum infection rates (MIR) in mosquitoes across Hungarian regions (NUTS-2), based on molecular xenomonitoring data (B). Regions: (1) Western Transdanubia, (2) Central Transdanubia, (3) Southern Transdanubia, (4) Central Hungary, (5) Southern Great Plain, (6) Northern Great Plain, and (7) Northern Hungary

### National temporal trends and disease-specific comparison

National monthly RSV values showed a significant temporal increasing trend over years (F = 58.3, p < 0.001). A significant Year × Search Term interaction was observed (F = 34.5, p < 0.001), indicating that temporal trends differed among the considered disease categories. Based on the interaction model, heartworm-related searches increased significantly over time (β = 1.78 ± 0.16 SE, p < 0.001). In contrast, the temporal slope for parvovirus was significantly smaller (β = −1.46 ± 0.23 SE, p < 0.001), and the slope for rabies was negative relative to heartworm terms (β = −1.75 ± 0.23 SE, p < 0.001), indicating no meaningful increase over time (Fig. 2). These findings indicate that the observed increase in search activity was specific to dirofilariosis and not attributable to a general rise in veterinary-related searches.

**Figure 2.**
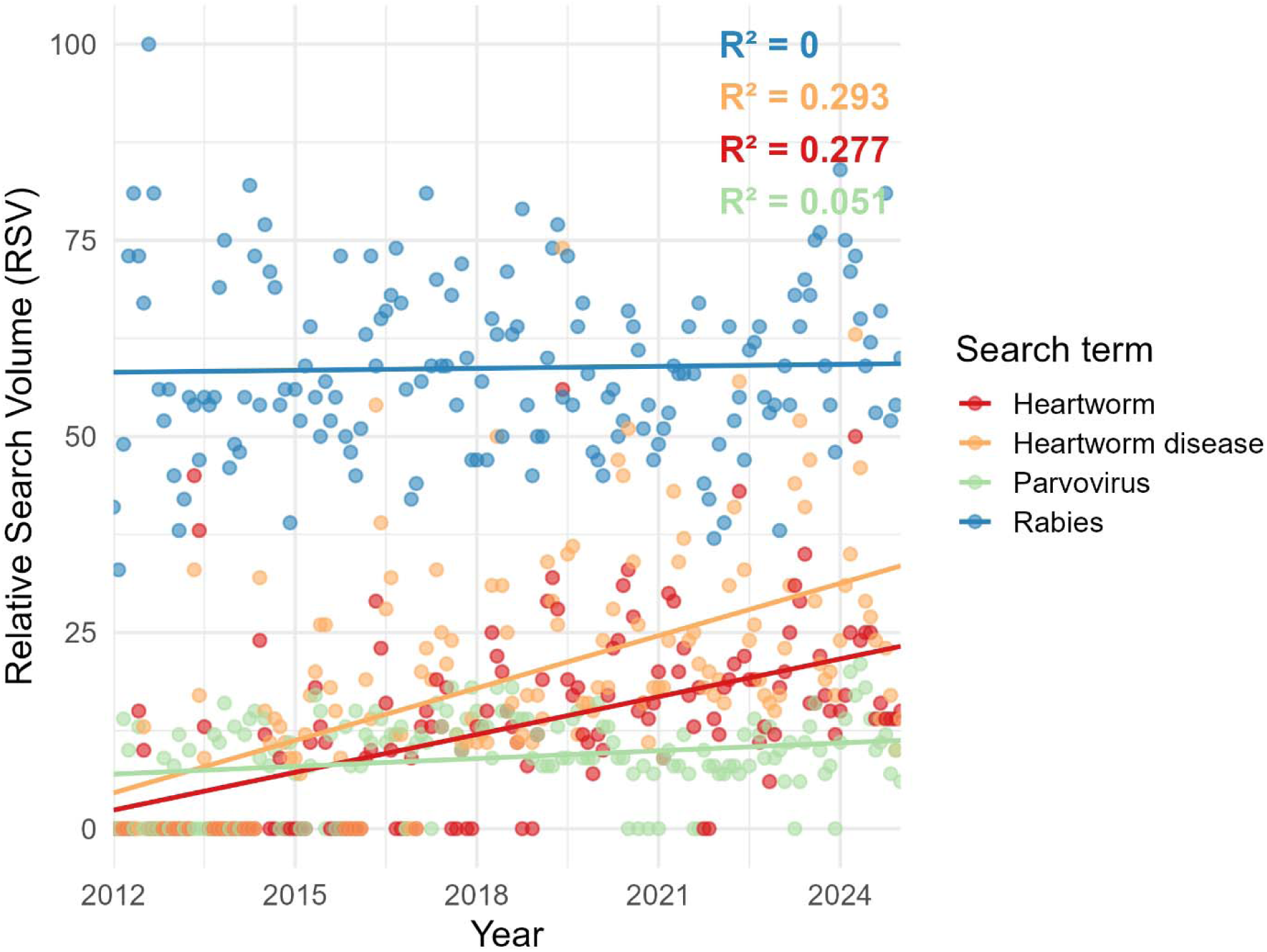
Temporal trends in Relative search volume (RSV) for canine diseases in Hungary from 2012 to 2025. Heartworm-related terms (heartworm as “szívféreg”, and heartworm disease as “szívférgesség”), parvovirus (“parvovírus”), and rabies (“veszettség”) are shown, with R2 calculated to each search term.

Significant seasonal variation in heartworm-related search activity was detected after accounting for long-term temporal trends (model comparison: F11,155 = 16.31, p < 0.001) (Fig. 3, Table 1). Monthly RSV values were highest between March and October, with peak search activity occurring during late spring and early summer (April–June), indicating seasonal patterns in public interest in heartworm disease, independent of the long-term increase in search activity over the study period.

**Figure 3.**
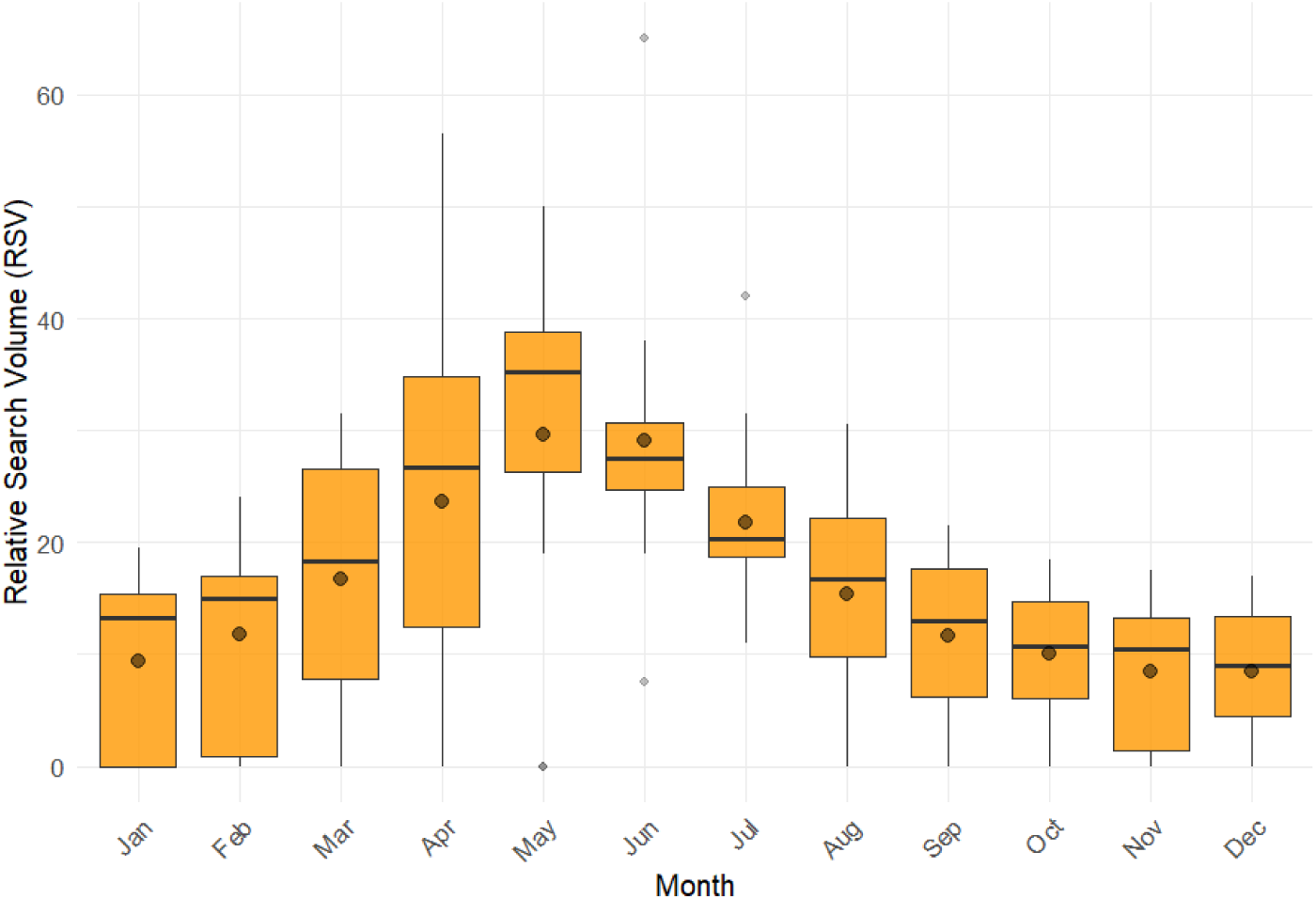
Seasonal variation in heartworm-related online search activity. Distribution of monthly relative search volume (RSV) values for heartworm-related search terms (average of “szívféreg” and “szívférgesség”) in Hungary over the period 2012–2025.

**Table 1.** Linear regression model evaluating seasonal variation in Google Trends relative search volume (RSV) for heartworm-related search terms (“szívféreg” and “szívférgesség”) in Hungary between 2012 and 2025. Monthly RSV values were averaged across the two heartworm-related search terms. January served as the reference month; positive coefficients indicate higher RSV relative to January after accounting for the long-term temporal trend.

| Predictor | Estimate ($\beta$) | SE | t value | p-value |
| --- | --- | --- | --- | --- |
| Intercept | -3574.70 | 280.94 | -12.72 | <0.001 |
| Year | 1.78 | 0.14 | 12.76 | <0.001 |
| February | 2.32 | 2.75 | 0.85 | 0.400 |
| March | 7.29 | 2.75 | 2.65 | 0.009 |
| April | 14.25 | 2.75 | 5.18 | <0.001 |
| May | 20.18 | 2.75 | 7.34 | <0.001 |
| June | 19.71 | 2.75 | 7.17 | <0.001 |
| July | 12.29 | 2.75 | 4.47 | <0.001 |
| August | 5.96 | 2.75 | 2.17 | 0.032 |
| September | 2.18 | 2.75 | 0.79 | 0.429 |
| October | 0.68 | 2.75 | 0.25 | 0.805 |
| November | -0.93 | 2.75 | -0.34 | 0.736 |
| December | -1.00 | 2.75 | -0.36 | 0.716 |

### Regional temporal patterns

Heartworm-related searches increased by an estimated 2.71 RSV units per year on the county scale (β = 2.71 ± 0.66 SE, t = 4.13, p < 0.001; Table 2), indicating an overall increase in heartworm-related search activity between 2012 and 2025. Allowing counties to vary in their temporal trajectories significantly improved model fit compared with a random-intercept model (χ² = 30.57, df = 2, p < 0.001), demonstrating that the rate of increase differed among counties. County-specific trends extracted from the random-slope model showed the strongest increases in Békés, Hajdú–Bihar, Heves, and Borsod counties (Fig. S1), whereas several western and central counties exhibited weak or non-significant temporal trends. Counties with the highest long-term mean RSV values were Csongrád (67.2), Jász-Nagykun-Szolnok (64.6), and Békés (50.6), indicating persistently elevated public interest in southeastern Hungary. Repeatability, calculated from the random-intercept model, was 0.39, indicating that 39% of the total variation in RSV was attributable to persistent differences among counties, while the remaining 61% reflected temporal variation within counties.

**Table 2.** Linear mixed-effects model evaluating temporal trends in county-level RSV values (2012–2025). RSV was modeled as a function of calendar year, with county included as a random intercept and random slope to account for repeated measurements and allow region-specific temporal trends.

| Component | Parameter | Estimate | SE | df | t value | p-value |
| --- | --- | --- | --- | --- | --- | --- |
| Fixed effects | Intercept | 7.58 | 5.10 | 19 | 1.49 | 0.154 |
| | Year ($\beta$) | 2.71 | 0.66 | 19 | 4.13 | < 0.001 |
| Random effects (County) | Intercept variance | 406.27 |  |  |  |  |
|  | Year slope variance | 6.65 |  |  |  |  |
|  | Intercept–slope correlation | −0.48 |  |  |  |  |
| Residual | Residual variance | 442.76 |  |  |  |  |

## Discussion

Our findings are consistent with previous studies demonstrating that Google Trends data can capture meaningful patterns of infectious disease activity. Most importantly, the strong positive correlation between regional Google Trends relative search volume (RSV) and mosquito minimum infection rates (MIR) provides independent support that online search activity reflects the known spatial distribution of *Dirofilaria* transmission in Hungary. Similar associations between online search intensity and epidemiological or entomological indicators have been reported for other vector-borne diseases, including West Nile virus and Lyme borreliosis in Europe and North America [33,52,53]. These findings suggest that online interest in heartworm-related search terms reflects the underlying ecological and epidemiological patterns of *Dirofilaria* transmission rather than random fluctuations in internet search behavior. Although search activity is inevitably influenced by factors such as veterinary communication, media coverage, and broader public discussion. This interpretation is further supported by veterinary records and molecular surveys documenting the expansion of canine dirofilariosis in Hungary [12–15,17], as well as by the absence of comparable temporal increases in the control search terms (rabies and parvovirus).

The seasonal peaks observed in GT data may support some biological relevance of search behavior. Most search activity occurred during late spring and summer, overlapping with the beggining of the active period of mosquito vectors. This seasonal co-occurence has been documented in other vector-borne diseases monitored via digital epidemiology, including West Nile virus and Lyme borreliosis, where search interest responds to vector abundance and seasonal risk perception [39,40]. Although dirofilariosis may take 6–9 months to produce clinical signs in dogs, patency can persist for up to 7.5 years in rare cases [54], with symptoms potentially appearing at any time of year in dogs. Therefore, seasonal increases in search activity are more likely to reflect routine veterinary surveillance and owner communication at the beginning of mosquito season rather than the onset of clinical disease. Heartworm screening is typically performed in early spring as mosquito activity begins, prior to initiating preventive treatment, which also increases the likelihood of detecting latent infections during this period.

The highest mosquito infection rates and search intensity were observed in southern and southeastern Hungary, regions repeatedly identified as hotspots for autochthonous transmission [12,15]. The geographic overlap between digital indicators and mosquito surveillance further supports the biological relevance of Google Trends as a proxy for regional *Dirofilaria* transmission. This combined approach shows the potential of GT as a complementary, low-cost surveillance tool for mapping emerging vector-borne diseases.

Nevertheless, some limitations must be highlighted. Regional RSV values may be less reliable in sparsely populated areas where search activity is low [55]. GT data also cannot directly measure disease incidence, clinical diagnoses, or vector infection prevalence [56]. Furthermore, our results demondtrate significant temporal increases and regional differences in search activity related to canine heartworm disease, which patterns may support interpretations about changes in distribution over time; however, our analysis did not directly assess disease emergence or spread. Importantly, while our analysis showed that regional search activity was strongly associated with regional mosquito infection rates, we did not directly test correlations between historical transmission data and longitudinal search trends. Therefore, the detected temporal patterns are consistent with documented epidemiological trends and may plausibly reflect underlying transmission dynamics, although this relationship was not directly evaluated in the present study. Nevertheless, the observed growth in search activity corresponds with increasing veterinary reports of canine infections and the expanding presence of competent mosquito vectors [12,13,59,14,15,17–19,51,57,58]. Overall, GT should be viewed as a supportive rather than standalone tool, particularly in settings where traditional veterinary epidemiological data are limited.

Despite these limitations, our findings are consistent with broader research demonstrating the value of digital epidemiology for complementing conventional infectious disease surveillance. For canine dirofilariosis, an expanding vector-borne disease of increasing veterinary and public health importance, Google Trends data may provide a useful supplementary source of information for identifying spatial and temporal patterns of disease activity, supporting retrospective analyses, and informing predictive models of disease distribution. Such information could help identify regions of increasing concern, guide targeted communication strategies, and support the allocation of resources for diagnostic testing and vector control. As climate change and invasive mosquito species continue to reshape the epidemiology of filarioid parasites across Europe, integrating digital behavioral data with entomological and veterinary surveillance could improve early detection and real-time awareness [60,61].

## Acknowledgements

Not applicable.

## Supplementary material

Data supporting findings and conclusions can be found in the manuscript, and in Supplementary Table 1 and Supplementary Figure 1.

